# Intravital optoacoustic ultrasound bio-microscopy reveals radiation-inhibited skull angiogenesis

**DOI:** 10.1101/500017

**Authors:** Héctor Estrada, Johannes Rebling, Wolfgang Sievert, Daniela Hladik, Urs Hofmann, Sven Gottschalk, Soile Tapio, Gabriele Multhoff, Daniel Razansky

**Author notes:** equal contribution (Héctor Estrada, Johannes Rebling).

## Abstract

Angiogenesis is critical in bone development and growth. Dense, large-scale, and multi-layered vascular networks formed by thin-walled sinusoidal vessels perfuse the plate bones and play an important role in bone repair. Yet, the intricate functional morphology of skull microvasculature remains poorly understood as it is difficult to visualize using existing intravital microscopy techniques. Here we introduced an intravital fully-transcranial imaging approach based on hybrid optoacoustic and ultrasound bio-microscopy, allowing for large-scale observations and quantitative analysis of the vascular morphology, angiogenesis, vessel remodeling, and subsurface roughness in murine skulls. Our approach also enabled high-throughput physiological studies to understand radiation-inhibited angiogenesis in the skull bone. We observed previously undocumented sinusoidal vascular networks spanning the entire skullcap, thus opening new vistas for studying the complex interactions between calvarian, pial, and cortical vascular systems.

## Main

The murine skullcap (calvaria) accommodates a multi-layered vascular network formed by sinusoidal vessels which are supplied by pial arteries and drained into dural veins (1–3). The calvarian vasculature is responsible for delivering oxygen and nutrients and impacts on bone marrow maintenance, development (4, 5), and hematopoiesis (6, 7), but can also be indicative of pathological changes upon radiotherapy (8). Despite its physiological importance, morphology and function of the calvarian vasculature remain poorly understood due to the lack of imaging tools capable of visualizing the large vascular network inside the optically scattering and highly heterogeneous bone plates (9). This gap in understanding is of particular significance with respect to radiation-induced vascular and bone marrow damages after targeted cytoreductive radiotherapy (RT), which has become a standard tool in clinical cancer treatment (10, 11). Despite significant progress in precise treatment targeting, collateral damage to neighboring tissue is unavoidable resulting in a number of significant side effects (12). Pathological changes to the parenchyma, the stroma, and the vasculature associated with long term cognitive disability have been observed in mammalian and human brains for single RT doses exceeding 10 Gy (13, 14). Likewise, RT treatment damages to the bone marrow microenvironment and vasculopathy have been reported in both long (5, 15) and plate (16) bones. In particular, calvarian plate bones contain a complex sinusoidal system, which collects newly formed cells from the bone marrow and releases them into the vascular system. The sinusoidal vessel walls are comprised of a single layer of endothelial cells supported by the surrounding bone marrow stroma (11). Due to their delicate structure, sinusoidal vessels are known to be strongly affected by ionizing radiation by inducing dilation, leakage, and necrosis (17, 18).

While tomographic imaging techniques can visualize larger vasculature in human skull (19), they lack the micrometer-scale resolution to resolve intricate sinusoidal networks (20). Histological techniques combined with cell-specific staining render high-resolution images of the vessels within the bone-marrow compartments of the skull, but are limited to observations in thin, *ex vivo* slices (5). Confocal and two-photon microscopy have been used to image flow dynamics (7), hematopoietic stem cell niche (6), and endothelial microdomains for tumor engraftment in bone marrow microvasculature (8). Intravital microscopy was applied to study histopathological changes in the murine bone marrow microvasculature following radiation therapy, revealing severe vascular damages (21). Optical microscopy approaches yet share two major drawbacks, namely, very restricted field-of-view and low penetration depth, both restricting imaging to small (sub-mm) regions of the skull and precluding investigations into the large-scale microvascular networks spanning the entire calvaria. More recently, optoacoustic (OA) microscopy has emerged as a powerful tool to image morphology and function of murine cerebral vasculature at multiple spatial scales (22, 23). By exploiting the strong optical absorption of hemoglobin in the visible spectral range, vascular networks can be imaged solely relying upon intrinsic contrast. Previous OA imaging studies focused on cerebral vasculature by using juvenile mice (four weeks old) where the skull tends to be significantly less vascularized, hence providing an unobstructed view of the underlying brain vasculature. In these studies, both the skull and the calvarian vasculature were therefore largely ignored (22) or considered a limiting factor (20).

Here, we present a new approach combining ultrasound (US) and OA bio-microscopy to visualize and precisely localize calvarian vasculature over the entire parietal and large parts of the frontal and interparietal skull bones. The high-resolution skull anatomy, retrieved from the automatically co-registered US data, is used for unequivocal differentiation between the cerebral and calvarian vasculature acquired by the OA modality. The method is applied to quantitatively evaluate radiation-inhibited angiogenesis in the murine skullcap.

## Results

### Intravital microscopy of radiation-induced vasculopathy

The hybrid OA and US bio-microscope for volumetric, label free intravital imaging (Fig. 1*A*, see Online Methods for details) targets the complex vascular network supplying the calvarian bone marrow (BM) compartments (Fig. 1*B*). Image-guided partial irradiation was performed on four-week-old female C57Bl/6 mice (n=7) using a small-animal radiation research platform (SARRP, X-Strahl, United Kingdom). Imaging was performed at the age of 15 weeks with the skull remaining intact and the scalp carefully removed. To generate mechanical contrast representing the three-dimensional skull bone morphology, our system utilized multiple US reflections resulting from a strong mismatch between elastic properties of skull bone and surrounding soft tissue. Fig. 1*C* thus reveals the calvarian bone plates, their sutures and major cerebral landmarks of the murine skull, such as the bregma and lambda (λ) junctions (24). Imaging of skull and brain vasculature was instead performed by focusing nanosecond laser pulses with a custom-designed gradient index (GRIN) lens and detecting the generated OA responses by the same transducer used for the US reflection-mode imaging. The low numerical aperture (NA=0.025) of the GRIN lens resulted in an extended depth-of-field (DOF), thus enabling three-dimensional imaging of large curved areas (see inset in Fig. 1 *C*) spanning most of the parietal and parts of the frontal and interparietal bones (Fig. 1*D*). As expected, calvarian bones, bone marrow, and the underlying cerebral cortex exhibit a densely vascularized and highly heterogeneous morphology (6). However, the sinusoidal vascular networks remained intact only in the non-radiated hemisphere. It should be noted that no fluorescence microscopy method was so far able to image such a large and curved FOVs in three dimensions. In addition, multiple fluorescent agents and imaging instruments are typically required to reliably differentiate between the skull and brain vasculature (2, 25). Imaging at different time points after irradiation (Fig.1*E* and *F*) reveals that the skull vasculature of the left hemisphere did not developed.

**Figure 1.**
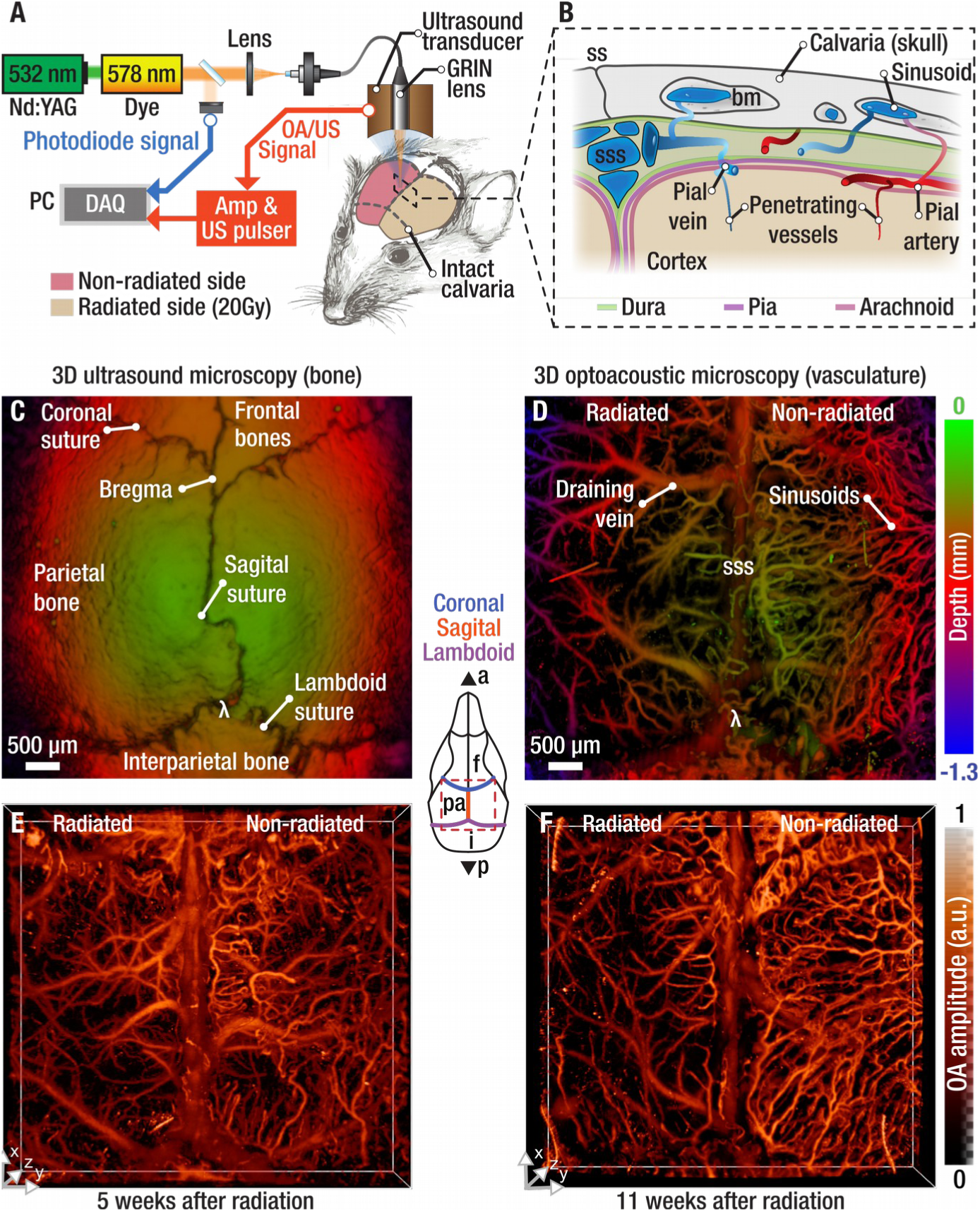
Label-free intravital ultrasound (US) and optoacoustic (OA) bio-microscopy of large-scale calvarian vascular networks in the murine skull. **A** Experimental setup utilizing a hybrid scan head to render fully co-registered volumetric images of the skull mechanical properties and vascularization. **B** Anatomy of the imaged vessels found in the murine calvaria and the underlying cortical meninges. **C** Depth-encoded maximum intensity projection of a volumetric US scan. **D** The simultaneously acquired volumetric OA image. **E** and **F** Maximum intensity projection (MIP) of the three-dimensional OA image acquired at 5 and 11 weeks after irradiation. Insert shows the approximate location of the 5.5 mm x 6.0 mm scanned area. ss – sagittal suture; sss – superior sagittal sinus; bm – bone marrow; a – anterior; f – frontal bone; pa – parietal bone; i – interparietal bone; p – posterior.

### Subsurface roughness analysis

Distinguishing cerebral from calvarian vasculature is challenging when solely relying on the OA data. This is because the OA-generated waves are distorted significantly when traversing the skull (26, 27), altering their shape, amplitude, and propagation velocity. The skull curvature further precludes unambiguous depth sectioning. We therefore developed an additional methodology (see Online Methods and Supplementary Fig. 4 for details) for accurate segmentation of the calvaria and extraction of its elastic and structural properties based on the naturally co-registered and depth-resolved OA (Fig. 2*A*) and reflection US (Fig. 2*B*) data. The depth-resolved segmentation of the vascular networks is facilitated by analyzing the pulse-echo waveforms to calculate the speed of sound in the skull using a homogenous solid plate model (Fig. 2 *C*). The combination of the OA and US waveforms in each point along the lateral dimensions subsequently reveals the actual location of the vasculature with respect to the skull (Fig. 2*D* and Supplementary Fig. 4).

**Figure 2.**
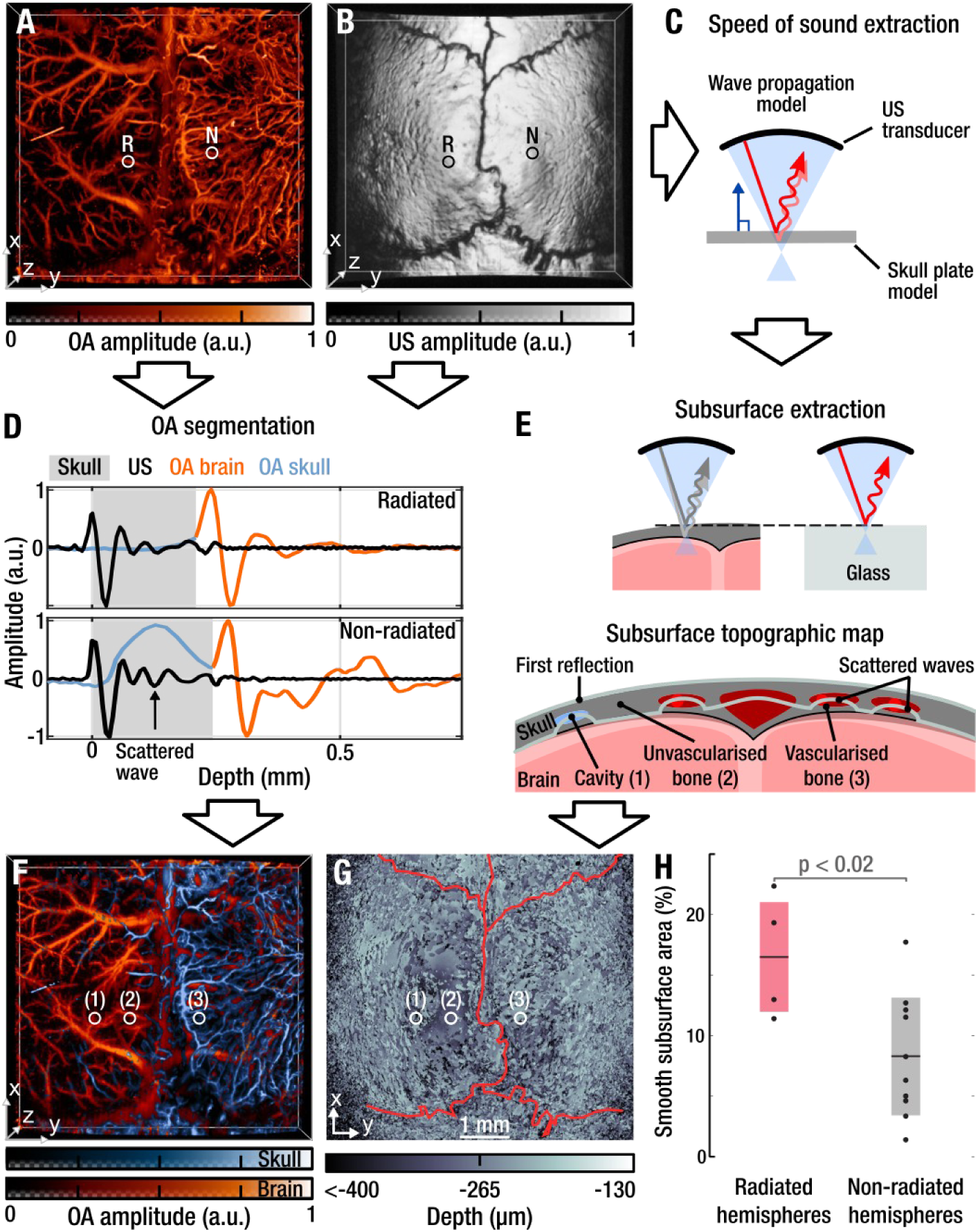
Accurate depth-resolved segmentation of the calvaria and extraction of its elastic and structural properties based on the naturally co-registered and depth-resolved OA and reflection US datasets. **A** Representative maximal intensity projection (MIP) of a volumetric OA dataset acquired from a partially radiated mouse. The radiated (I) and non-radiated (N) hemispheres are labeled with circles. **B** The simultaneously acquired three-dimensional pulse-echo US image. **C** The pulse-echo waveforms are used to extract the speed of sound using a solid plate model (see Online Methods and Supplementary Fig. 5 for details). Combination of the OA and US waveforms, obtained from the points labeled in **A** and **B**, reveals the actual location of the vasculature with respect to the skull **D**, which allows the segmentation in skull (blue) and brain (orange) vasculature **F**. Scattering events in the US data **D** can be used, after cross correlation with a measured glass reference **E**, to inspect the bone morphology below its surface **G**, where the color scale represents the apparent distance from a given scattering event to the skull surface. **H** Statistical analysis on seven mice (four radiated, three sham-radiated) shows a larger portion of smooth (roughness < 14 µm) subsurface skull topography in the radiated hemispheres (see also Supplementary Fig. 4 and Supplementary Table 1). Black dots - individual mice, black lines – mean value, grey and red bars – standard deviation. One-way ANOVA F test yields p=0.01885.

The segmented depth-resolved map of the vascular networks (Fig. 2*F*) clearly reveals the absence of calvarian vasculature in the radiated hemisphere. The voids associated with the vasculature and bone marrow compartments in the skull are responsible for generating additional US scattering (Fig. 2*E*). This can be shown by the presence of additional peaks following the main reflection peak at the skull’s surface (Fig. 2*D*). By means of a glass plate reference and cross correlation to disregard spurious peaks, we obtained a topographic map of the skull’s subsurface (Fig. 2*G*), which is indicative of the skull’s heterogeneities. In contrast to the non-radiated hemisphere (Fig. 2*G*), the radiated hemisphere contains relatively smooth areas close to the sagittal suture (Fig. 2*G* (2)), in agreement with the segmented OA image showing no vasculature in those areas. A higher degree of roughness is manifested in other non-vascularized areas (Fig. 2*G* (1)), which could be ascribed to voids left by inactive vasculature, bone marrow, or other structural bone inhomogeneities. Statistical analysis of skull’s subsurface roughness in partially radiated (n=4) and sham-radiated (n=3) mice (Fig. 2*H*, Supplementary Fig. 6, and Supplementary Table 1) shows a significantly increased area of non-vascularized bone in the radiated hemispheres (17% mean) as compared to sham-radiated controls (8% mean).

### Large-scale imaging of calvarian vasculature

We were able to visualize the distinct and intricate vascular networks found in the murine calvaria over regions exceeding 6 mm across, attaining spatial resolution of 12 µm and 30 µm in the lateral (x, y) and axial (z) dimensions, respectively (Fig. 3). It is noteworthy that segmentation of skull vasculature (blue) from the underlying cortical vessels (red) relies entirely on the endogenous contrast of bone and blood in the US and OA modes. Figs. 3*A*–*C* show the segmented and intact calvarian vasculature found in the sham-radiated mice. All calvarian bones contain bone marrow microenvironments (28) which is evident from the strong vascularization present in the frontal, parietal, and interparietal bones. Previous studies have found highly branched and densely connected sinusoids in the diaphysis of long bones (5), a growth pattern that can also be observed in the parietal bone vessels in Figs. 3*A*–*C*. Unlike previously reported (8), calvarian vasculature was not only located parasagittally within the frontal and parietal bones but was instead present across the entire parietal bones in the sham-radiated mice. The area around the coronal suture is also highly vascularized, in agreement with previous findings by Mazo *et al.* (9), where large veins located in the frontal bone (arrow heads in Figs. 3*A*–*C*) connect to sinusoidal networks propagating in parallel to the irregularly shaped coronal suture. The large draining veins in the frontal bones are partially linked to the similarly sized vessels oriented in parallel and slightly lateral to the sagittal suture, where they presumably drain into the superior sagittal sinus of the brain. In contrast, the lambdoid suture exhibits sparse vascularization (arrows in Figs. 3*A*–*C*), despite the known presence of large bone marrow compartments in the neighboring interparietal bone (28). The parasagittally located veins in the parietal bone form an irregularly-shaped sinusoidal network characterized by a highly variable pattern of multiple interconnecting side branches. The sinusoids within the parietal bones have relatively constant diameter. This pattern significantly differs from the cerebral vasculature which normally has a better-defined hierarchy of small venules connecting into larger draining veins, eventually terminating into the superior sagittal sinus.

**Figure 3.**
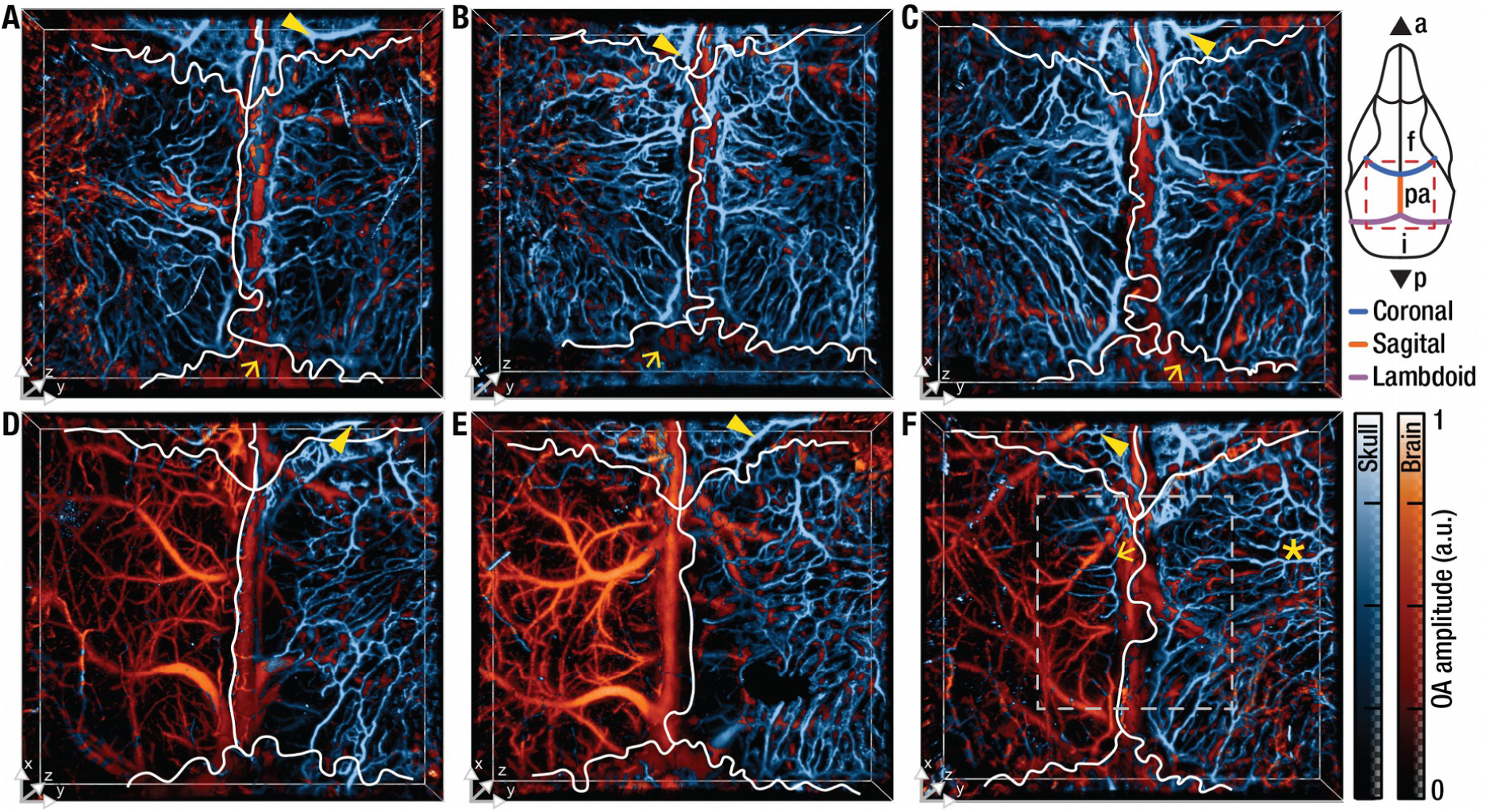
Volumetric segmentation of microvasculature in radiated and sham-radiated mice. **A-C** Representative examples of the volumetric optoacoustic data segmented for the skull (blue) and brain (orange) vasculature in sham-radiated mice; **D**-**F** The corresponding images acquired eleven weeks post irradiation. White lines correspond to the skull’s sutures extracted from ultrasound data (see Supplementary Fig. 4). a – anterior; f – frontal bone; pa – parietal bone; i – interparietal bone; p – posterior.

### Vasculopathy following partial radiation exposure

We subsequently employed the hybrid microscopy system to study the effects of targeted RT on the large-scale calvarian vasculature. To this end, the left hemisphere of four-week-old female C57Bl/6 mice (n=4) was radiated with a mean dose of 20 Gy (see online methods). Imaging of the calvarian microvasculature was performed 11 weeks following the partial irradiation treatment. The non-radiated, right hemispheres in Figs. 3*D*–*F* show a dense network of sinusoidal vessels similar to those found in the sham-radiated controls in Figs. 3*A*–*C*. In stark contrast, the calvarian microvasculature is nearly missing in the (radiated) left hemispheres thus exposing the underlying cerebral vasculature. This is in agreement with previous reports which identified significant regression of the sinusoidal endothelium and substantial decline in endothelial cells after radiation exposure (11). The dense vascular network located near the coronal suture in the sham-radiated mice is missing in all but one radiated mouse where some vasculature is still present (Fig. 3*F*). A number of small vessels are also present in the frontal bones in all the radiated mice (arrow heads in Figs 3*D*–*F*), but their number and density is strongly reduced compared to the sham-radiated controls while the larger draining veins are mostly missing. The area around the sagittal suture in the parietal bones is similarly devoid of vascularization, again with the exception of one mouse in which a single thin vessel branches laterally and terminates at ∼1 mm distance (see arrow in Fig. 3*F*). Notably, the large parasagittal draining veins found laterally to the sagittal suture in both parietal bones of the sham-radiated mice are largely missing in the radiated mice. This is the case not only for their radiated left hemisphere but also for the right hemisphere, despite precise targeting of the irradiation treatment (see Online Methods). Instead, the dense sinusoidal network of the right hemisphere drains into areas of the parietal bone located more laterally (see asterisk in Fig. 3*F*).

### Quantitative vasculopathy analysis

To quantify the effects of the targeted irradiation on the calvarian vasculature, we developed an automatic vessel segmentation and analysis (AVSA) algorithm (see online methods and Supplementary Fig. 7). In short, maximum intensity projections of the calvarian vasculature along the depth dimension are filtered and thresholded and input into the vessel characterization algorithm, which accurately tracks the individual vessel segments and assesses their diameter and branch points (Fig. 4*A*, *B*). Using AVSA, we separately analyzed the segmented calvarian vascular network of left and right hemispheres of both radiated and sham-radiated mice. The retrieved vascular parameters were subsequently compared using a one-way analysis of variance (ANOVA) where p-values <0.05 were considered statistically significant. Quantification of the microvasculature in the calvaria of the sham-radiated animals found an average of 364 and 328 vessels for the left and right hemispheres, respectively (Fig. 4*C*). A similar number of 315 vessels was found in the non-radiated, right hemisphere of the radiated mice. In contrast, vessel numbers significantly diminished in the left hemisphere of the mice where an average of only 82 vessels was found (p<0.001), corresponding to a 77% reduction. Likewise, the number of branch points detected in the vascular network was significantly reduced in the radiated hemisphere (n=24) as compared to left (n=171) and right (n= 144) hemispheres of the sham-radiated mice as well as the non-radiated right hemisphere (n=145, all p<0.001), i.e. 84% vessel reduction (Fig. 4*D*). The more significant decline of the branching points as compared to the vessel numbers indicates a diminished complexity of the remaining vascular network with less vessels branching and looping. A less drastic trend was observed when analyzing the average length of the individual vessel segments (Fig. 3*E*). The vessels in the right hemisphere of sham-radiated control mice (180 µm) and non-radiated right hemisphere of the targeted mice (171 µm) were significantly longer than those in the radiated left hemisphere (143 µm, p=0.009 and p=0.003 respectively). A similar trend is observed for the diameter of the vessel segments (Fig. 4*F*) where the radiated hemisphere contained significantly smaller vessels (42.7 µm average vessel diameter) as compared to the non-radiated hemisphere (50.2 µm, p = 0.003) and the right hemisphere of sham-radiated mice (50.9 µm, p=0.003). The intact parts of the radiated vascular network show a less significant decline of 18% and 14% in vessel length and diameter, respectively, suggesting a partial preservation of the calvarian vasculature, as can be also observed in Figs. 2*F* and 3*F*.

**Figure 4.**
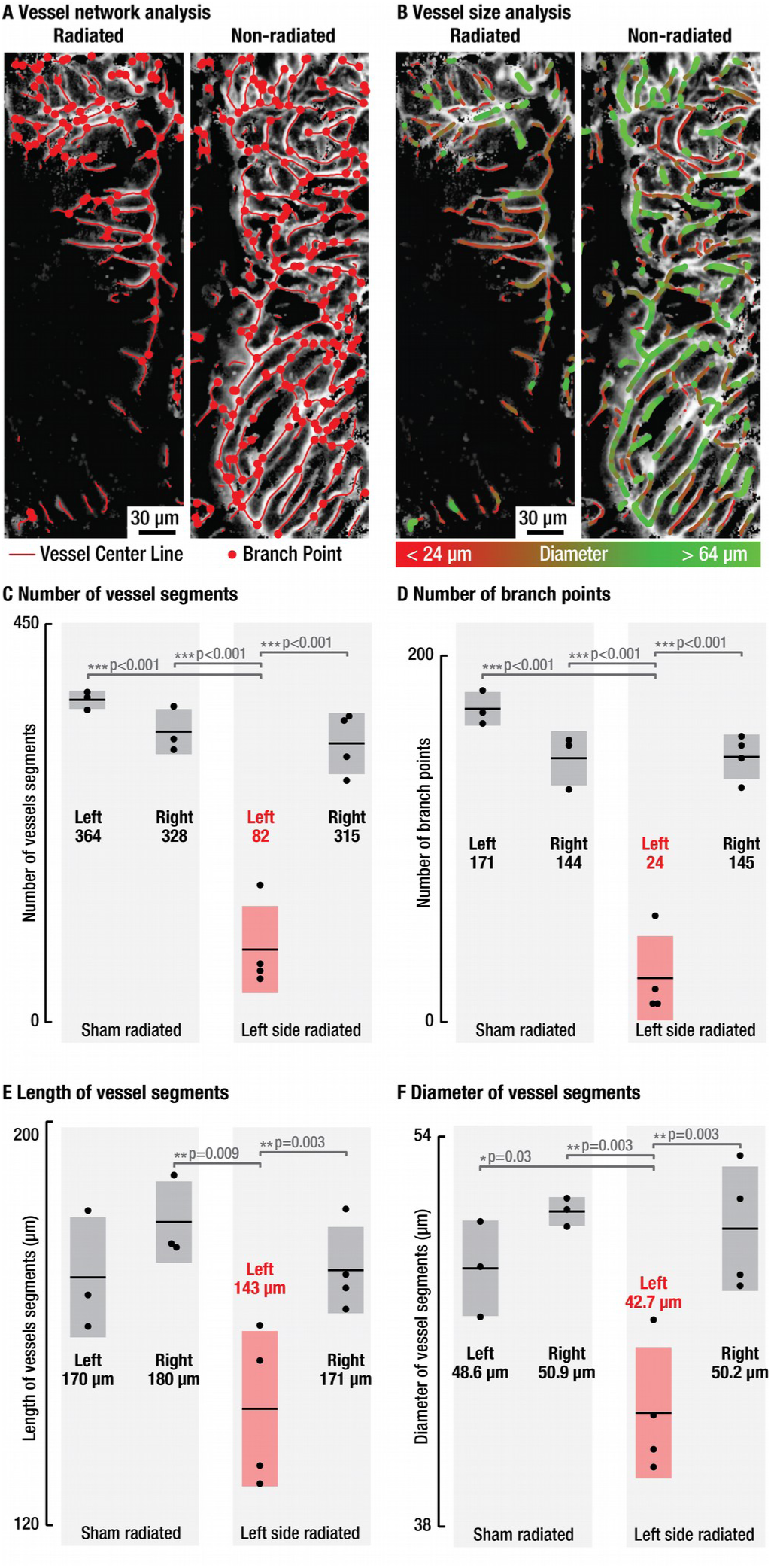
Quantitative vessel network analysis reveals significant loss in calvarian sinusoids following targeted irradiation. The calvarian vasculature in the segmented optoacoustic datasets served as the basis for automatic detection of vessel positions and branching points **A** as well as vessel diameters **B**. The image area used to illustrate the quantitative vasculature analysis is indicated by gray dashed line in Fig. 4F. For the statistical analysis, all segmented skull vessels datasets were analyzed. A strong and statistically significant decline in the number of individual vessels segments (**C**, 77% reduction) and branching points (**D**, 84% reduction) was induced by the targeted irradiation. In contrast, both the length (**E**) and diameter (**F**) of vessels that resisted the irradiation treatment was reduced by only 18% and 14%, respectively. Black dots - individual mice, black lines – mean value, grey and red bars – standard deviation.

## Discussion

The bone marrow located in the calvarian plate bones is an important site for the generation and preservation of blood and immune cells and is maintained by a multilayered microvasculature network, the calvarian sinusoids (29). The latter provide an interface between the haematopoietically active bone marrow and the peripheral circulation and form an important niche for hematopoietic progenitor differentiation (11, 16) whose function is currently poorly understood. Previous investigations have looked into the bone-marrow vascular niche function in murine femurs using slow and invasive histological techniques (5) or otherwise were limited to small areas of the calvaria further requiring administration of exogenous contrast agents (6).

Our study is the first to observe intact bone marrow microvasculature over a centimeter-scale FOV spanning the entire skullcap, further eliminating the need for surgical alteration and vascular labeling due the strong absorption contrast of hemoglobin. We further implemented a synergistic combination of the concurrently acquired volumetric US and OA data to unequivocally segment the calvarian vasculature from the underlying cerebral vasculature. The intact vascular network was imaged without the need for tiling or depth-scanning, thus enabling large-scale observations of the vascular morphology, angiogenesis, and vessel remodeling. In this way, we were able to unveil a complex and dense vascular network present in the parietal plate bone of the skull. While calvarian vasculature has recently come into the research spotlight (5, 7, 21), its functional anatomy remains under substantial debate (2). In contradistinction to previous reports where bone marrow microvasculature was only found near the coronal and sagittal sutures (6), we observed a highly heterogeneous vascular anatomy with a dense network of sinusoids spanning the entire bone plate, particularly in the parietal areas.

The delicate vessels spanning most of the calvaria are known to be strongly affected by ionizing radiation (10). This has important implications for targeted tumor treatments as well as radiative conditioning prior to bone marrow transplantation (12). In our study, targeted radiation treatment of one hemisphere resulted in a significant supression of vasculature growth over the entire radiated hemisphere observed at 5 and 11 weeks following the procedure. This effect was revealed both by the stand-alone US modality as well as via the combined OA and US bio-microscopy approach. The structural information of the skull bone retrieved by the US data alone is not sufficient to accurately assess the vascular network. Nonetheless, the extracted subsurface topology allows inspecting the skull’s subsurface roughness to identify any unvascularized areas in a completely noninvasive manner since US imaging does not require scalp removal. The inhibited angiogenesis is also readily apparent in the segmented OA images as all the radiated parietal bones were nearly devoid of vasculature, while the underlying brain vasculature appeared intact. This effect could be attributed to the delicate structure of the sinusoidal vessels, which, unlike the cerebral veins, consist of a single thin layer of endothelial cells (11). The rapidly self-renewing endothelial cells are particularly sensitive to irradiation insults where damage may appear weeks or even months following the procedure (14). The transcranial imaging capability thus enables physiological long-term studies crucial for understanding the radiation-inhibited angiogenesis and vessel remodeling. Elucidating the dose dependence and time course of calvarian vascular damage may facilitate optimal targeting of RT doses, thus resulting in better treatment outcomes.

The bone marrow sinusoids are presently believed to recover from the radiation exposure via angiogenesis rather than neovascularization (16). This implies that the vascular damage is chiefly repaired by division of the existing sinusoidal endothelial cells, already part of the vascular wall, rather than proliferation of new endothelial cells. Our study showed no evidence of angiogenesis or recovery in the radiated hemisphere 11 weeks post 20 Gy partial radiation treatment. It therefore appears that the precursors of endothelial cells were damaged irreversibly so the endothelial layer could no longer be created via angiogenesis, resulting in a complete breakdown of the vascular network. We additionally observed a decline of calvarian vasculature near the sagittal suture in the non-targeted hemispheres. Such loss of larger draining vessels can be ascribed to an abscopal effect of the irradiation that was otherwise carefully targeted to a single hemisphere. Beyond the field of radiation treatments, the newly developed approach opens new venues for systematic high-throughput studies of the murine calvarian vasculature. In humans, a striking variance in the distribution and density of skull vasculature among different individuals has been previously reported (30). However, no comparable large-scale investigations into the murine calvarian vasculature were so far possible. Our fast and scalable imaging setup enables the investigation of microvascularization across both highly localized as well as large areas of the skull, involving different animal strains and developmental stages. Knowledge of the developing calvarian vessels aids studies looking at subjects exhibiting strong vascularization with rich bone-marrow microenvironment. Furthermore, animals exhibiting sparsely developed sinusoidal networks can be used to avoid potential sources of bleeding when applying intravital microscopy through cranial windows or visualizing cerebral vasculature with OA microscopy, where calvarian vasculature may partially obstruct the underlying brain vessels. Finally, the transcranial large-scale US and OA bio-microscopy approach could also aid in more efficient guidance of localized fluorescence microscopy studies looking at vascular features (21) or stem cell homing (31). Further developments of our technique could enable *in vivo* studies of the complex interactions between calvarian, pial, and cortical vasculature over a large portion of the murine skull.

## Acknowledgements

We gratefully acknowledge the technical assistance of Michael Reiss during the imaging experiments. This work was supported in part by the European Research Council grant ERC-2015-CoG-682379 and the German Research Foundation (DFG) Grant RA1848/5-1.

## Author Contributions

JR and DH conceived the study; WS performed the radiation experiments; HE, WS and DH performed the imaging experiments; HE, JR and UH processed the image data; HE and UH developed the vessel segmentation algorithm and subsurface skull roughness analysis; JR developed the AVSA algorithm and performed statistical analysis; DR, ST and GM supervised the study; All authors contributed to the design of experiments, discussing the results and writing the manuscript.

## Methods

No statistical methods were used to predetermine sample size. The experiments were not randomized and investigators were not blinded during experiments and outcome assessment.

### Imaging Setup

The experimental setup used throughout this study is schematically depicted in Fig. 1. Imaging was performed by a custom-designed hybrid OA and US bio-microscopy system, whose detailed optical, mechanical, and electrical designs are illustrated in Supplementary Fig. 1. Central to the microscope’s design is a fast-moving scan head sampling the specimen data in reflection mode. The OA and US imaging modalities share the same scanning mechanism and scan head, resulting in inherently co-registered volumetric OA and US datasets. This simplifies the subsequent post-processing steps as no software-based image registration is required.

A Q-switched diode-pumped Nd:YAG laser (IS8II-E, EdgeWave, Germany) pumping a dye laser (Credo, Sirah Lasertechnik, Germany) tuned to a 578 nm wavelength were used for the OA signal excitation. The laser generates 10 ns duration pulses with a pulse repetition frequency of up to 10 kHz. The energy of the dye laser output can be freely adjusted using a half-wave plate (AHWP10M-600, Thorlabs, USA) in combination with a polarizing beam splitter (PBS251, Thorlabs, USA).

The 578 nm excitation light is subsequently coupled into a large-mode-area photonic crystal fiber (PCF, model: LMA-20, NKT Photonics, Denmark). The PCF is endlessly single-mode, has a large effective mode field area of approximately 215 µm², and low losses to allow high power delivery without nonlinear effects or material damage. Efficient coupling into the low NA (0.02) PCF was achieved by focusing the 4 mm diameter laser beam through a 100 mm focal distance lens (AL50100M, Thorlabs, USA). Coupling efficiency above 20% was accomplished by mounting the proximal end of the PCF onto a 6-axis kinematic mount (K6XS, Thorlabs, USA) and by careful alignment at low laser energies to avoid damaging the fiber facet. The distal end of the PCF terminates in a gradient-index (GRIN) lens (GRINTECH, Jena, Germany) mounted in a 1 mm diameter central aperture of a spherically-focused polyvinylidene fluoride US transducer (Precision Acoustics, Dorchester, United Kingdom), as illustrated in Supplementary Fig. 1. The GRIN lens focuses the excitation light onto a diffraction-limited 12 µm spot at a distance of 6.5 mm. The low numerical aperture of the lens (0.025) results in an extended depth of focus (DOF) of at least 2mm, as measured by a *√ 2* factor broadening of the spot size. The axial resolution (along the depth dimension) of about 30 µm is mainly determined by the 30 MHz effective bandwidth of the US detector (30 MHz central frequency) while the lateral acoustic resolution in the pulse-echo imaging mode was 45 µm and 20 µm axially (Supplementary Fig. 2). The optical focus of the GRIN lens and the acoustic focus of the US transducer are coaxially and confocally aligned, eliminating the need for an optical-acoustic beam combiner (22) and resulting in an optimized detection sensitivity. The high sensitivity in combination with the extended DOF enabled OA imaging of the entire murine skull without the need for averaging, depth scanning, or image tiling.

During volumetric image acquisition, the scan-head is rapidly scanned in a zig-zag pattern in the xy-plane by a combination of a fast linear stage (M-683, PI, Germany) and a slower stage (LTM 60F-25 HSM, OWIS, Germany). In OA imaging mode, the detected OA signals are recorded by the US transducer and amplified by an 8 dB pre-amplifier (Precision Acoustics, United Kingdom) and a 24 dB broadband low-noise amplifier (ZFL-500LN, Mini-Circuits, USA). For each laser pulse, the analog signals are digitized at 14 bit using a two-channel 250 MS/s data acquisition (DAQ) card (M3i.4142, Spectrum Systementwicklung Microelectronic, Germany). The per-pulse laser energy is simultaneously monitored by sampling 1% of the laser’s output using a beam sampler (BSF10-A, Thorlabs, Newton, USA) positioned in front of the PCF. The sampled beam was measured by a fast, calibrated photodiode (DET10A, Thorlabs, USA) and digitized by the second channel of the DAQ. Pulse-echo US imaging was performed using a pulser-receiver (5073PR, Olympus, Massachusetts, USA) for generation of broadband US pulses and the same DAQ card for sampling the reflected time-resolved signals. The digital raw data recorded during the imaging was buffered in memory and was saved for post-processing. The system control was executed by a custom software written in Matlab 2017b (Mathworks, United States) and C++ (via Matlab MEX files). Data recording and post-processing were performed on a personal computer using an Intel Core i7-6800k processor clocked at 3.40 GHz with 64 GB memory.

### Animal experiments

All *in vivo* animal experiments were performed in full compliance with the institutional guidelines of the Helmholtz Center Munich and the Technical University of Munich and with approval from the Government District of Upper Bavaria. Image-guided partial irradiation was performed on four-week-old female C57Bl/6 mice (Charles River, United States) using a small-animal radiation research platform (SARRP, X-Strahl, United Kingdom). Mice were anaesthetized by isoflurane/oxygen inhalation for the duration of the procedure. Transverse, sagittal and frontal computed tomography scans using 60 kV and 0.8 mA photons were performed for each mouse for precise radiation targeting (32). The irradiation targeted 40% of the left brain volume with a beam size of approximately (6×8) mm^2^ and with a mean target dose of 20 Gy using 220 kV and a 13 mA X-ray beam filtered with copper (0.15 mm). The software SARRP control and Muriplan (both X-Strahl, United Kingdom) were used for CT imaging, precise radiation targeting, and calculation of the irradiation dose. A total of four mice received partial brain irradiation while three mice where sham radiated. All mice fully recovered after the procedure and were housed in single ventilated cages under pathogen-free conditions until imaging.

### Intravital imaging

Imaging of the cerebral and calvarian microvasculature was performed 5 and 11 weeks after partial irradiation treatment. The mouse body temperature was monitored and maintained using a rectal thermometer coupled to a feedback-controlled heating pad. Additionally, blood oxygenation and heart rate were continuously monitored throughout the experiments (PhysioSuite, Kent Scientific, USA). The anesthetized mice were head-fixed in a custom-designed mouse head holder (Narishige International Limited, UK) to avoid motion artifacts. During the measurement the mice were kept anesthetized using isoflurane (1% - 2% v/v) in 100% O_2_ supplied via a breathing mask integrated into the mouse head holder. The scalp was incised in the midline using micro-dissecting scissors (Sigma-Aldrich, Germany) to expose the parietal bones and parts of the frontal and interparietal bones without damaging the calvaria. The exposed calvaria was cleaned using hemostatic sponges (Gelfoam®, Pfizer, United States) to remove remaining hair and stop any bleeding. A thin US gel layer was applied to the calvaria to provide acoustic coupling and maintain the skull hydrated. A water-filled Petri dish having a 2 cm opening covered with a thin polyvinylidene chloride foil was attached to the mouse head with the microscope’s scan-head submerged into water in order to allow unperturbed acoustic signal propagation (see Supplementary Fig. 1). All images were obtained without introduction of exogenous contrast agents. The maximal laser power on the specimen during OA imaging was 7 mW (1 µJ per-pulse energy, 7 kHz pulse-repetition rate, 10 ns pulse duration). Recording of raw data across a FOV of 6.0 mm x 5.5 mm took approximately two minutes for OA imaging and less than one minute for US imaging. All mice were sacrificed under anesthesia immediately following the imaging experiment.

### Data post-processing

The unfiltered US and OA datasets are stored during the imaging session and post-processed to create volumetric representations of the imaged area. For this, the raw data are band-pass filtered using a first order Butterworth filter between 1 MHz and 60 MHz to match the frequency response of the US transducer (from around 15 to 45 MHz, −6dB bandwidth) and cancel out DC components in the digitized signals without losing information. The OA signals are further corrected for laser power fluctuations using the simultaneously recorded photodiode signals. To compensate for irregularly spaced data created by the continuous zigzag scanning pattern, we applied linear scattered data interpolation to the OA and nearest-neighbor interpolation to the US data due to the different lateral resolution and smoothness between both imaging modalities. Each time-resolved A-scan (single point of the xy-scan) was encoded with an imaging depth using the speed of sound in water of 1481 m/s. All post-processing, including vessel segmentation and analysis was carried out using Matlab 2017b (Mathworks, United States) and Python 2.7.12 (Python Software Foundation). The post processing of raw data took approximately 30 s per dataset. Visualization of three-dimensional data was performed using Mayavi2 4.4.3 (33).

### Ultrasound data processing for vessel segmentation

Due to the strong mismatch in the elastic properties between bone and soft tissue, the high amplitude signal reflected from the skull can be cross correlated with a pulse-echo reference signal recorded from a flat glass reference in order to map the skull surface. The pulse-echo reference was measured for different axial locations in steps of 10 µm and constitutes and experimental characterization of the electro-acoustic properties of the US transducer when interacting with a fluid-solid interface (34).

For a given A-scan, the glass reference with the same (or closest) axial location is selected to perform the cross correlation, whose maximum is used to calculate the skull’s surface depth. Repeating this process for all the scanned points yields the outer skull surface. The skull’s subsurface roughness is subsequently calculated by extracting the positive peaks of the cross correlation function and estimating their position relative to the outer skull surface, i.e.,

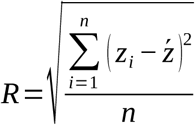

where *n* and *z_i_* denote the number of peaks within the region of interest and their distance from the skull and 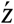 is the average distance.

One of the secondary cross correlation peaks corresponds to a reflection at the inner skull surface, but its classification depends strongly on the skull’s morphological, geometrical, and elastic properties. A few fiduciary points with clear US and OA features (see Supplementary Fig.4) were selected to guide a manual segmentation assuming a homogeneous thickness for each hemisphere. For simplicity, only the longitudinal speed of sound in the skull bone (3695 m/s) has been considered, which was estimated using an approximated physical model (20).

### Automatic vessel segmentation and analysis

En face images of skull and brain vasculature were generated by calculating maximum intensity projections (MIPs) of the segmented volumetric OA data along the depth dimension. The average number of detected vessel segments in these MIPs for the imaged mice was 1041. To analyze this relatively large number of vessels we employed an automatic vessel segmentation and analysis (AVSA) algorithm that provides the precise number, position, and length of the individual vessel segments as well as the number and location of branch points (see Supplementary Fig. 7). Our analysis is derived from an algorithm originally developed for use on retinal images where it was previously shown to accurately measure various vessel parameters (35). The OA signals have a large dynamic range governed by the presence of both large, e.g. draining veins in the brain, and finer vessels, such as arterioles in the brain and vasculature in the skull. The large dynamic range is first compressed using contrast-limited adaptive histogram, providing an optimized image contrast. Next, the vessels are segmented from surrounding tissue by thresholding the contrast enhanced MIPs using a multiscale vessel enhancement filter (36). A number of simplified vessel detection algorithms have been proposed to extract parameters from the segmented binary images, but results depended strongly on the particular thresholding approach and have shown to be unreliable (35). In contrast, the AVSA algorithm computes the raw centerlines of potential vessels found in the binary image. Spline fitting is then used on the individual vessel segments to provide smooth and accurate vessel orientation and position. Next, a second derivative of the image pixel intensities is calculated perpendicular to the fitted vessels positions. A zero-crossing of the derivative is used to accurately identify the vessel edges on both sides of the centerline and the vessel diameter. It is noteworthy that the precise positions of the vessel segments (start point, end point, centerline) represent various morphologically relevant vascular parameters, such as vessel length, tortuosity, or branching angles. As the vessel segment positions are also known in a global coordinate system, additional network parameters, such as vessel diameter ratios during branching, vessel occupancy, and density, can be readily measured. We additionally analyzed the fractal dimension using the thresholded images. The entire vessel analysis procedure, including filtering, thresholding, and single vessel analysis, took approximately 10 s per single dataset.

### Statistical analysis

A number of key morphological parameters of both skull and vasculature were further analyzed to investigate the effects of the partial irradiation treatment more elaborately. Statistical analysis of the skull’s subsurface roughness is based on the differences in roughness distribution (see Supplementary Figure 6) between radiated and non-radiated hemispheres. After setting a roughness threshold at 14 µm, we calculate the relative area of smooth subsurface (< 14 µm) for each hemisphere and perform a one-way ANOVA F test (Supplementary Table 1) using Python 2.7.12 (Python Software Foundation). Statistical analysis of the vascular network in the presented study encompassed the vessel area density (number of vessels per mm^2^), number of the reliably identified individual vessel segments, number of branch points, length of the vessel segments, vessel diameter and fractal dimension. For each mouse, the OA datasets were divided into left and right hemispheres based on the location of the sagittal suture. Vessel analysis was carried out individually for the resulting 14 datasets (both hemispheres in n=7 subjects). The average value found in each dataset was used to statistically compare vessel segment diameters and lengths. As a result, statistical analysis was carried out using six data points for sham-radiated mice (both hemispheres in 3 mice) and using eight data points for radiated mice (both hemisphere in 4 mice). Statistical analysis was performed in Matlab 2017b (Mathworks, United States) using the build-in one-way analysis of variance (ANOVA). P-values were calculated using a F-test, where p-values < 0.05 were considered statistically significant. Boxplots in Fig. 3 show mean value (black line) and standard deviation (shaded box) while black dots denote values in the individual datasets. Detailed statistical results are shown in Supplementary Table 1 and 2.

