## Supplemental Figures and Tables for "Intravital optoacoustic ultrasound bio-microscopy reveals radiation-inhibited skull angiogenesis"

### Intravital optoacoustic microscopy reveals radiation-induced skull vasculopathy

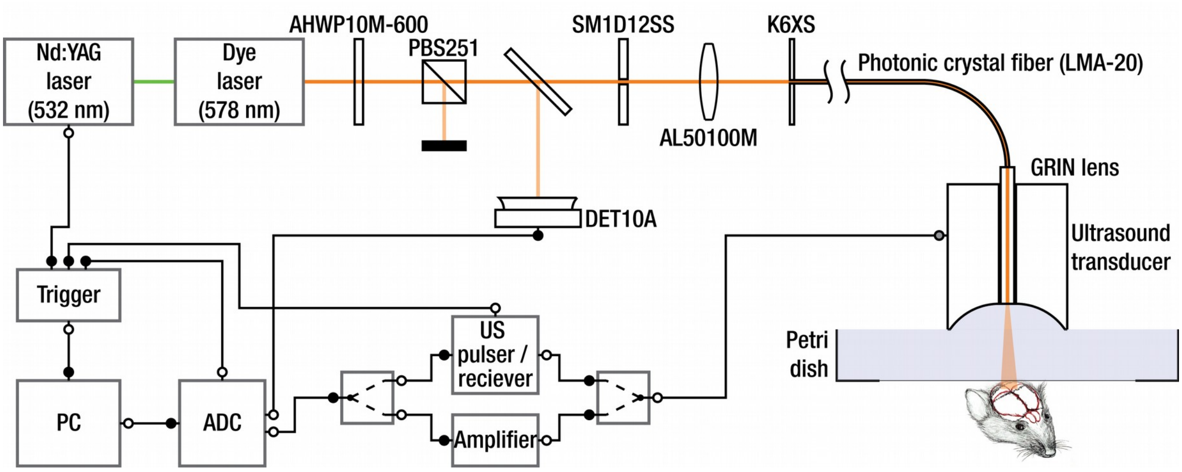


**Supplementary Figure 1. Detailed overview of the hybrid optoacoustic (OA) and ultrasound (US) bio-microscopy system.** Nanosecond-duration laser pulses (~1 µJ per-pulse energy) from a dye laser tuned to 578 nm and pumped by a 532 nm Nd:YAG laser are used to generated OA responses from the imaged tissue. Pulse energies are continuously monitored via the DET10A photodiode. Pulses are coupled into a photonic crystal fiber and focused onto the specimen using a gradient index (GRIN) lens. The OA signals generated inside the specimen are recorded by a spherically focused US transducer, amplified, and digitized. A pulser/receiver is used to generates short US pulses, which are subsequently focused into the specimen by the same transducer, which is also used to record the signals reflected from the tissue.

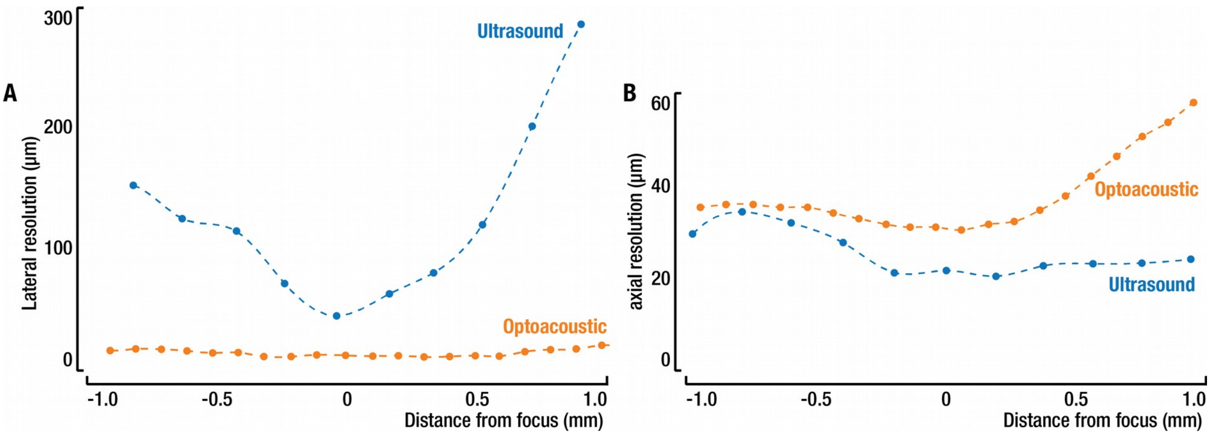


**Supplementary Figure 2. Resolution of hybrid optoacoustic (OA) and ultrasound (US) biomicroscope. A** Lateral and **B** axial resolution for the two imaging modalities as a function of the distance to the focus. A sharp silicon edge was used as target. The lateral resolution corresponds to the full width at half maximum (FWHM) of the line spread function for OA and US. The axial resolution for OA is obtained as the FWHM of the first peak and the FWHM of the pulse envelope (using Hilbert transform) for US.

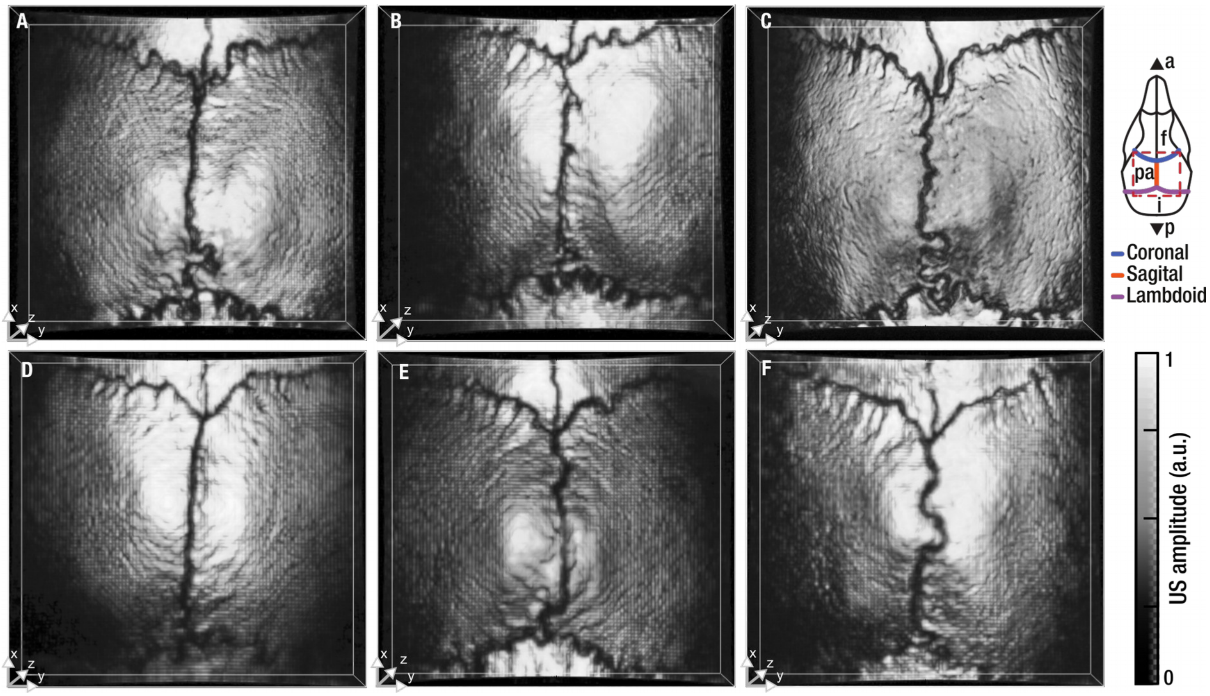


**Supplementary Figure 3. Ultrasound volumetric images of irradiated and sham-irradiated mice.** Maximum amplitude projection of the pulse-echo ultrasound data are shown. **A-C** sham -irradiated mice and **D-F** mice irradiated on the left hemisphere.

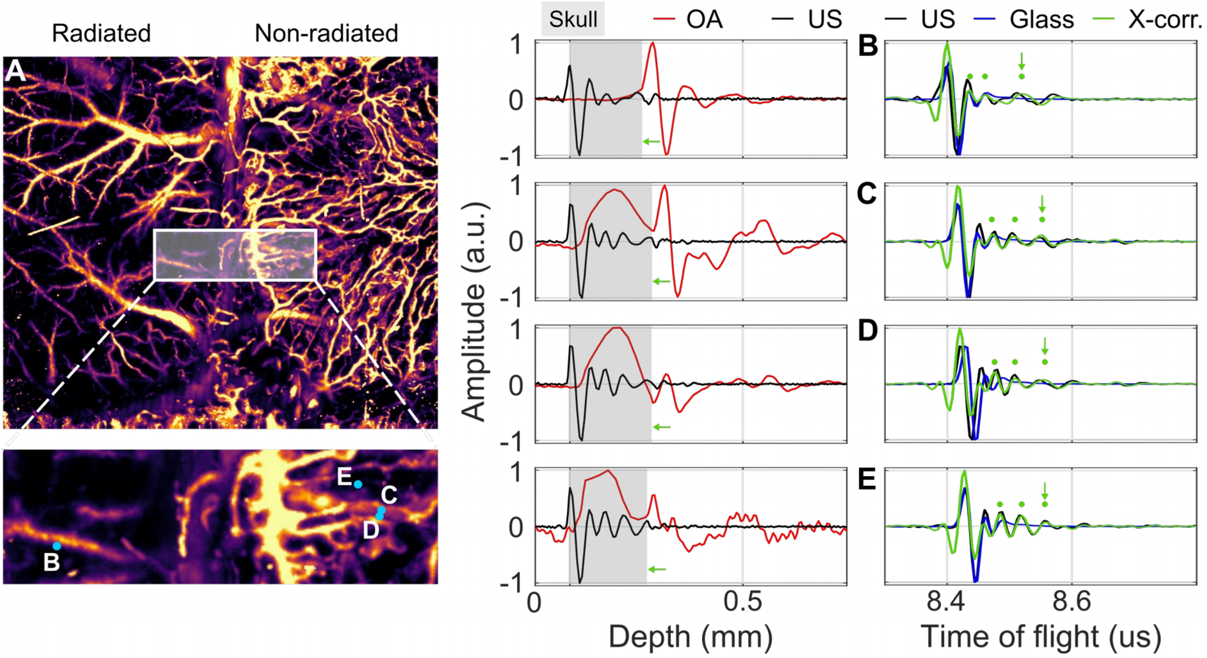


**Supplementary Figure 4. Combined optoacoustic (OA) and ultrasound (US) waveform analysis. A** Maximum amplitude projection (MAP) of the scanned mouse head region. The inset indicates zoomed-in region where positions of the analyzed points are labeled. Individual OA and US waveforms recorded in the different points - **B** Pial vasculature, **C** calvarian and pial vasculature, **D** calvarian vasculature, and **E** weak amplitude point localized in the skull, presumably bone marrow. Gray region corresponds to the skull location. Pulse-echo waveforms of the skull, glass reference, and their cross correlation are show in the right column. Green points show the position of the cross-correlation peaks whereas arrows point the inner skull surface.

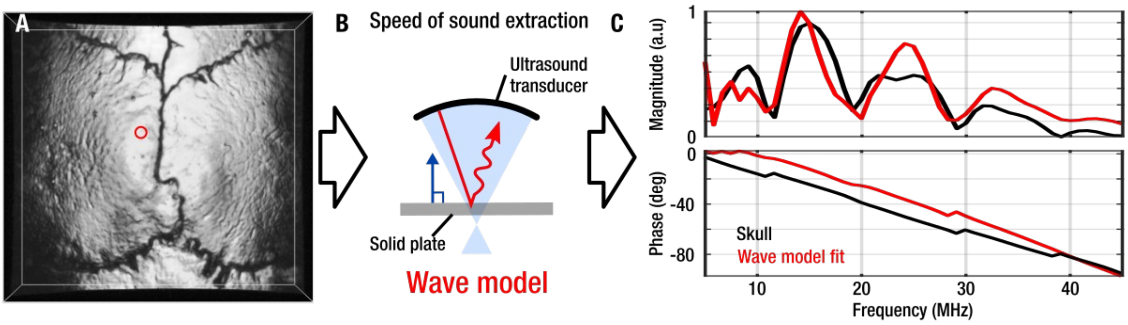


**Supplementary Figure 5. Extraction of the speed of sound from pulse-echo data. A** Maximum amplitude projection of the ultrasound reflection. A point (red circle) where the skull is perpendicular to the spherically focused transducer axis (for simplicity of the model) is converted to frequency domain using the Fourier transform. **B** A forward frequency domain model of a solid viscoelastic plate is solved and the elastic constants used as parameters subject to genetic algorithm optimization. **C** Comparison between the fitted model and the measurement in the skull.

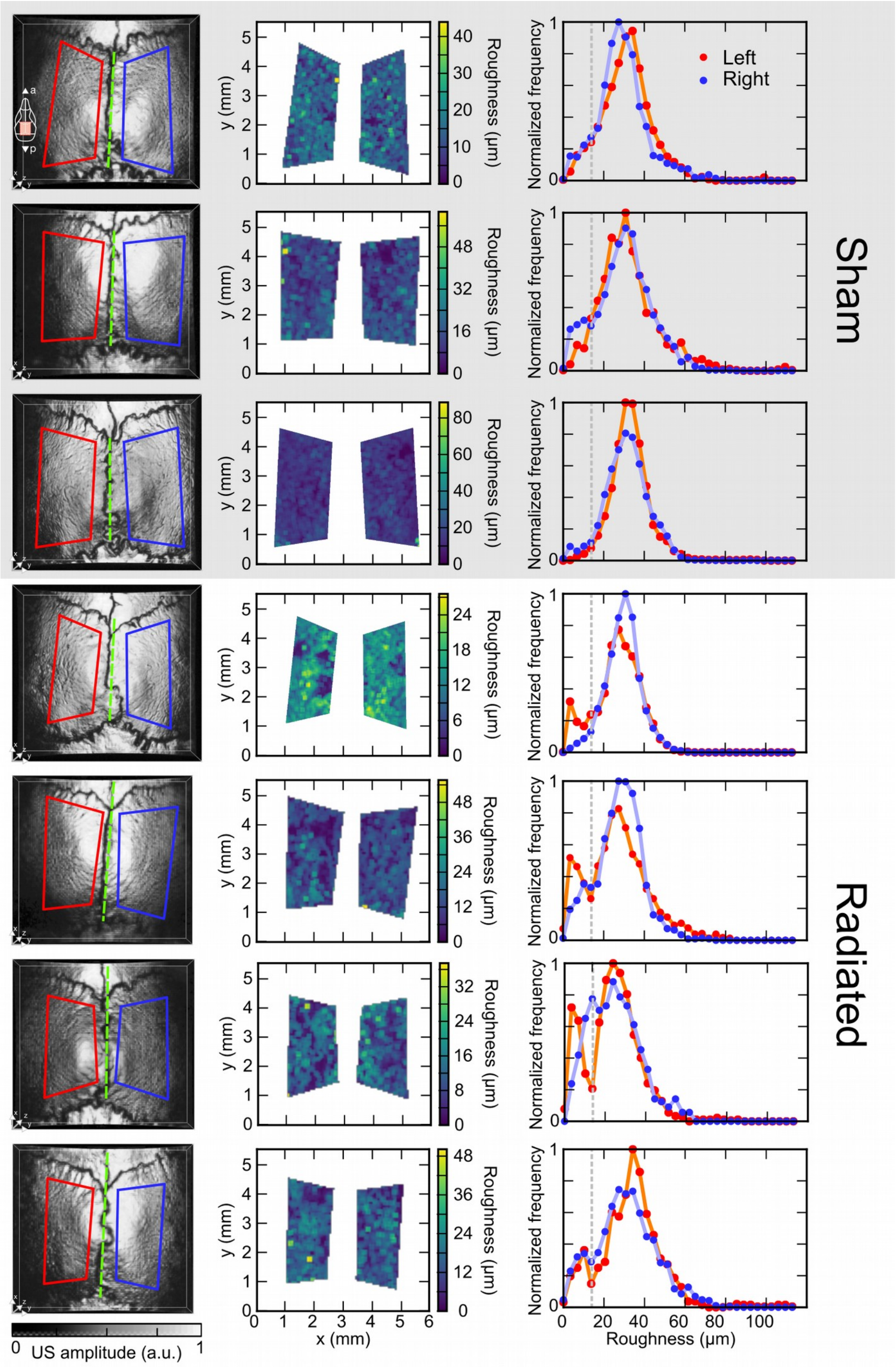


**Supplementary Figure 6. Skull's subsurface roughness analysis. A** The analyzed regions (red and blue trapezoids) and the symmetry axis (green dashed line) are indicated in the volumetric pulse-echo ultrasound data shown in the left column. **B** The skull subsurface roughness calculated using a 75 µm window on the 4th cross correlation peak (eight peaks detected in total). **C** Normalized frequency distribution of the skull's subsurface roughness for the different hemispheres. The dashed line indicates the 14 µm threshold.

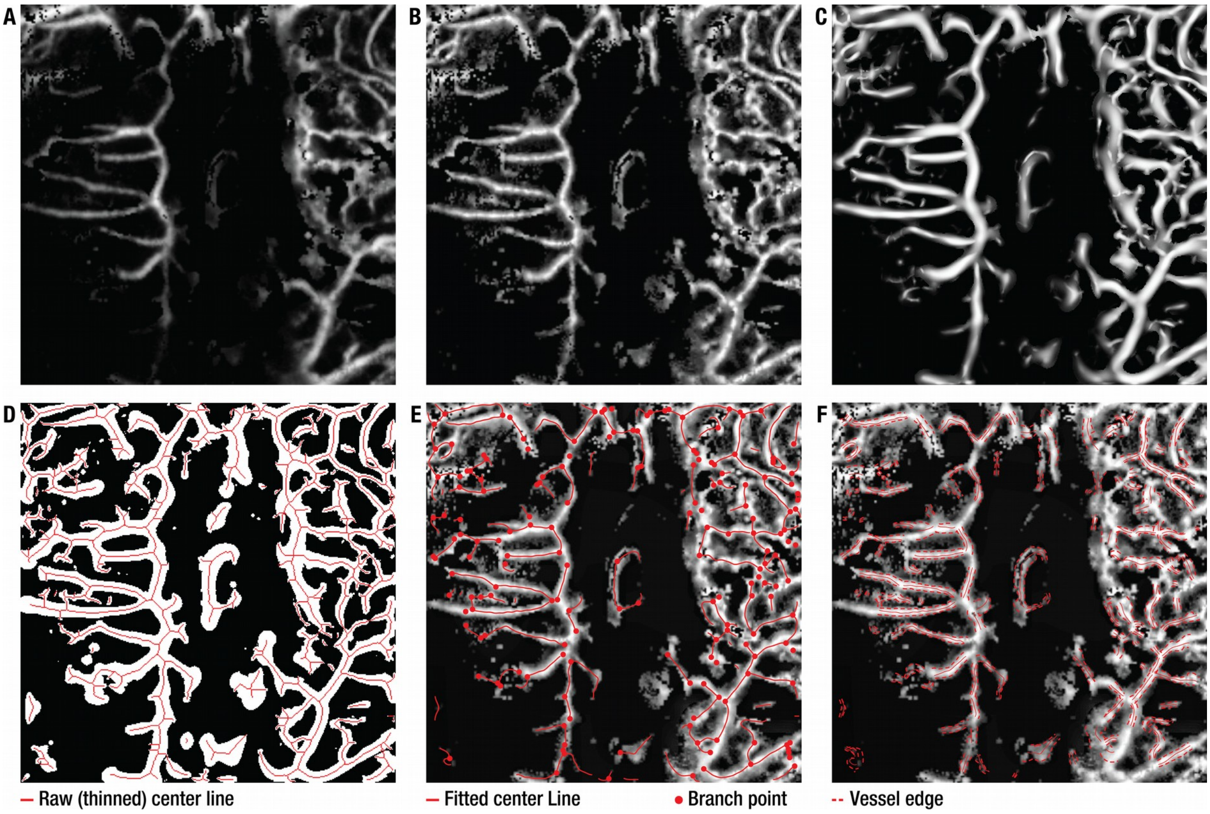


**Supplementary Figure 7. Automatic vessel segmentation and analysis (AVSA) algorithm. A** Raw optoacoustic MIP of the skull vasculature. **B** Contrast-enhanced MIP using contrast-limited adaptive histogram equalization. **C** Vasculature enhanced following the vessel-filtering. **D** The vessel-filtered image is segmented and reduced to raw vessel centerlines (red lines). **E** Centerlines found in **D** are the basis for spline-fitting of centerlines and branch detection. **F** The fitted centerlines are used to determine the vessel edges (red dashed lines) and hence the diameters perpendicular to the fitted centerlines.

**Supplementary Table 1. Overview of the extracted skull parameters for all mice.**

| Mouse | Hemisphere | Irradiated | Area with surface skull roughness < 14 μm [%] | Roughness dist. diff. between hemispheres [%] | Area with sub-surface skull roughness < 14 μm [%] | Assumed skull thickness [μm] |
|---|---|---|---|---|---|---|
| 1 | L | y | 32.1 | 14.2 | 19.3 | 203.6 |
| 1 | R | n | 24.1 | | 11.5 | 278.1 |
| 2 | L | y | 23.6 | 14.8 | 22.4 | 211.0 |
| 2 | R | n | 7.9 | | 17.7 | 298.0 |
| 3 | L | y | 12.1 | 11.9 | 11.4 | 223.5 |
| 3 | R | n | 38.5 | | 12.7 | 273.1 |
| 4 | L | y | 34.8 | 12.2 | 13.0 | 206.1 |
| 4 | R | n | 37.1 | | 3.4 | 285.5 |
| 5 | L | n | 25.0 | 13.0 | 6.3 | 260.7 |
| 5 | R | n | 42.6 | | 8.3 | 260.7 |
| 6 | L | n | 16.5 | 11.0 | 5.0 | 260.7 |
| 6 | R | n | 40.2 | | 12.1 | 260.7 |
| 7 | L | n | 12.6 | 9.9 | 1.4 | 260.7 |
| 7 | R | n | 28.3 | | 4.6 | 260.7 |

**Supplementary Table 2. Overview of the extracted vessel parameters for all mice.**

| Mouse | Hemisphere | Irradiated | Vessel length (µm) Mean | Std | Median | Min | Max | Vessel diameter (µm) Mean | Std | Median | Min | Max | Tortuosity Mean | Std | Median | Min | Max |
|---|---|---|---|---|---|---|---|---|---|---|---|---|---|---|---|---|---|
| 1 | L | y | 128.2 | 105.9 | 89.7 | 10.7 | 521.3 | 41.2 | 13.6 | 38.2 | 23.1 | 100.2 | 1.15 | 0.84 | 1.02 | 1.00 | 7.43 |
| 1 | R | n | 182.7 | 169.5 | 123.1 | 9.6 | 1295.8 | 47.9 | 15.7 | 44.5 | 19.2 | 124.5 | 1.06 | 0.12 | 1.02 | 1.00 | 2.17 |
| 2 | L | y | 159.6 | 144.9 | 90.0 | 9.8 | 804.8 | 40.4 | 9.8 | 38.4 | 25.4 | 83.4 | 1.06 | 0.13 | 1.02 | 1.00 | 1.65 |
| 2 | R | n | 167.0 | 170.5 | 100.7 | 9.9 | 1247.6 | 53.2 | 19.6 | 48.6 | 24.9 | 161.1 | 1.05 | 0.11 | 1.02 | 1.00 | 2.26 |
| 3 | L | y | 131.8 | 103.3 | 98.4 | 9.8 | 576.8 | 42.6 | 15.4 | 39.4 | 24.4 | 146.7 | 1.05 | 0.09 | 1.02 | 1.00 | 1.62 |
| 3 | R | n | 169.7 | 154.7 | 113.9 | 10.1 | 851.0 | 48.3 | 15.6 | 45.2 | 23.4 | 149.2 | 1.06 | 0.14 | 1.02 | 1.00 | 2.30 |
| 4 | L | y | 152.7 | 137.8 | 101.5 | 20.4 | 641.5 | 46.5 | 17.6 | 44.4 | 23.8 | 129.4 | 1.09 | 0.24 | 1.03 | 1.00 | 2.53 |
| 4 | R | n | 162.8 | 150.4 | 112.7 | 10.0 | 1337.9 | 51.5 | 18.3 | 46.5 | 24.4 | 159.4 | 1.05 | 0.10 | 1.02 | 1.00 | 2.09 |
| 5 | L | n | 165.6 | 149.8 | 109.3 | 9.5 | 802.2 | 46.6 | 16.6 | 43.6 | 19.5 | 164.8 | 1.07 | 0.20 | 1.02 | 1.00 | 3.93 |
| 5 | R | n | 175.8 | 186.7 | 111.0 | 10.3 | 1413.9 | 51.5 | 19.4 | 46.4 | 25.7 | 187.8 | 1.05 | 0.10 | 1.02 | 1.00 | 1.92 |
| 6 | L | n | 159.4 | 164.2 | 102.4 | 9.9 | 1279.8 | 48.7 | 15.8 | 46.0 | 24.1 | 161.3 | 1.06 | 0.16 | 1.02 | 1.00 | 3.56 |
| 6 | R | n | 175.1 | 163.2 | 122.7 | 9.9 | 1125.9 | 51.0 | 16.9 | 47.7 | 26.2 | 175.8 | 1.05 | 0.09 | 1.02 | 1.00 | 1.96 |
| 7 | L | n | 182.4 | 200.2 | 119.9 | 9.9 | 1953.2 | 50.5 | 19.9 | 46.5 | 18.8 | 155.3 | 1.06 | 0.13 | 1.02 | 1.00 | 2.15 |
| 7 | R | n | 189.4 | 183.4 | 124.5 | 10.1 | 1108.2 | 50.3 | 15.0 | 47.4 | 24.4 | 154.0 | 1.05 | 0.09 | 1.02 | 1.00 | 1.63 |

X

| Mouse | Hemisphere | Irradiated | Number of vessels [-] | Number of Branches [-] | Total Length (mm) [-] | Vessel Coverage (%) [-] | Vessel Area Density (vessels/mm²) [-] | Fractal dimension value | lower conf. | upper conf. |
|---|---|---|---|---|---|---|---|---|---|---|
| 1 | L | y | 58 | 10 | 7.4 | 2.6 | 3.2 | 1.29 | 1.22 | 1.36 |
| 1 | R | n | 273 | 128 | 49.9 | 20.5 | 18.4 | 1.54 | 1.49 | 1.58 |
| 2 | L | y | 49 | 10 | 7.8 | 2.8 | 3.0 | 1.29 | 1.23 | 1.36 |
| 2 | R | n | 300 | 144 | 50.1 | 18.9 | 18.2 | 1.53 | 1.49 | 1.57 |
| 3 | L | y | 155 | 58 | 20.4 | 7.7 | 9.5 | 1.41 | 1.35 | 1.46 |
| 3 | R | n | 341 | 151 | 57.9 | 20.9 | 20.7 | 1.52 | 1.48 | 1.56 |
| 4 | L | y | 66 | 18 | 10.1 | 4.4 | 4.0 | 1.39 | 1.33 | 1.45 |
| 4 | R | n | 346 | 156 | 56.3 | 21.9 | 21.0 | 1.54 | 1.49 | 1.58 |
| 5 | L | n | 367 | 163 | 60.8 | 21.8 | 22.4 | 1.51 | 1.47 | 1.55 |
| 5 | R | n | 320 | 151 | 56.3 | 21.4 | 19.4 | 1.56 | 1.52 | 1.61 |
| 6 | L | n | 373 | 181 | 59.5 | 22.5 | 22.7 | 1.51 | 1.47 | 1.55 |
| 6 | R | n | 357 | 154 | 62.5 | 23.8 | 21.7 | 1.55 | 1.51 | 1.59 |
| 7 | L | n | 353 | 169 | 64.4 | 23.9 | 21.5 | 1.54 | 1.50 | 1.57 |
| 7 | R | n | 308 | 127 | 58.3 | 21.2 | 18.7 | 1.55 | 1.51 | 1.59 |

X
